# Toxic stress-specific cytoprotective responses regulate learned behavioral decisions in *C. elegans*

**DOI:** 10.1101/2020.02.24.962381

**Authors:** Gábor Hajdú, Eszter Gecse, István Taisz, István Móra, Csaba Sőti

## Abstract

**Background:** Protection of organismal integrity involve physiological stress responses and behavioral defenses. Recent studies in the roundworm *Caenorhabditis elegans* have shown that pathogen and toxin exposure simultaneously stimulate cellular stress and detoxification responses and aversive behavior. However, whether a coordinate regulation exists between cellular and neurobehavioral defenses remains unclear.

**Results:** Here we show that exposure of *C. elegans* to high concentrations of naturally attractive food-derived odors, benzaldehyde and diacetyl, induces toxicity and aversive behavior. Benzaldehyde preconditioning activates systemic cytoprotective stress responses involving DAF-16/FOXO, SKN-1/Nrf and Hsp90 in somatic cells, which confer behavioral tolerance to benzaldehyde and cross-tolerance to the structurally similar methyl-salicylate, but not to the structurally unrelated diacetyl. In contrast, diacetyl preconditioning augments diacetyl avoidance and does not induce apparent molecular defenses. Reinforcement of the experiences using massed training forms relevant associative memories. Memory retrieval by the odor olfactory cues leads to avoidance of food contaminated by diacetyl and context-dependent behavioral decision to avoid benzaldehyde only if there is an alternative, food-indicative odor.

**Conclusions:** Our findings reveal a regulatory link between physiological stress responses and learned behavior which facilitates self-protection in real and anticipated stresses. The potential conservation of this somato-neuronal connection might have relevance in maladaptive avoidant human behaviors.

## Background

Adequate, coordinated responses of multicellular organisms are key to adapt to and overcome fundamental alterations of the environment (*1–3*). These responses originate from intracellular molecular defenses, such as the oxidative, xenobiotic, metabolic and proteotoxic stress responses, which guard homeostasis and confer cytoprotection against the respective stresses, promoting physiological adaptation, fitness and longevity at the organismal level (*4*). Adaptation also involve complex behavioral responses orchestrated by the neuroendocrine system (*5–7*). For instance, sensory cues representing danger evoke aversive behavior as a result of perception of multiple sensory stimuli, neuronal processing and decision making both in humans and in other species (*8–10*). In some cases, the neural impulse of perceived danger is so intense that the organism decides to avoid co-occurring cues representing life-sustaining qualities such as food (*6, 11*). Besides external sensory cues, decision making is modulated by neural context like arousal, motivation, and reward (*12, 13*). Importantly, behavioral decisions are also influenced by sensory cues that evoke associative memories of past events (*14*). Moreover, exaggerated, inadequate avoidant behavior is characteristic to human anxiety disorders such as phobias (*11*), where sometimes intense physical symptoms of toxicity and disgust are evoked by olfactory cues. Although the neuroendocrine mechanisms of stress are extensively studied, the contribution of somatic, especially intracellular defenses to behavioral regulation is largely unknown.

The soil nematode *Caenorhabditis elegans* with its 959 cells is a versatile model system to study the link between cytoprotective stress responses and behavior. Worms, using a well-defined network of 302 neurons are capable of complex behavioral decisions (*15–17*). Flavors and volatiles have great impact on decision making of nematodes, informing about possible nutrition and danger *via* neuronal processing of olfactory and gustatory cues (*16*). Besides well characterized escape responses, tissue damaging insults, such as toxins and pathogens, induce a network of evolutionary conserved cytoprotective defenses in each somatic cell and in specialized tissues (*4*). Fixing the actual damage as well as eliminating damaging agents are key mechanisms of cellular protection (*18*). Nematodes and mammals share diverse molecular processes to recognize and overcome toxic, stressor agents, such as the FOXO and Nrf2 pathways. A key oxidative and metabolic stress response regulator in *C. elegans* is the FOXO ortholog DAF-16 transcription factor (*19*). DAF-16 is ubiquitously expressed, localized in the cytosol, and is activated by nuclear translocation in response to oxidative and genotoxic agents, starvation, desiccation and heat stress (*20*). Loss-of-function mutations or RNAi knockdown of *daf-16* results in compromised resistance to multiple stresses and shorter lifespan (*21*).

The Nrf-2 orthologue SKN-1 transcription factor is the major xenobiotic and oxidative stress regulator in nematodes (*18, 22*). Its nuclear translocation is induced by dietary restriction, pathogen attack, the INS/IGF-1 and TIR-1/PMK-1 pathways to modulate cellular respiration, enhance oxidative stress resistance, immunity and systemic detoxification defenses (*22–24*). SKN-1 cooperates with numerous stress related pathways and regulators including DAF-16 and the *C. elegans* heat-shock transcription factor orthologue HSF-1 to fine-tune cytoprotective gene expression patterns (*18*). Upregulation of specific and overlapping molecular stress responses underlies an adaptive process called physiological conditioning hormesis in stress biology (*25*). In course of hormesis, a conditioning stress exposure results in increased survival under a subsequent, lethal stress evoked by the same or a different stressor, a phenomenon called stress tolerance or cross tolerance, respectively. To clearly discriminate physiological and behavioral terms, we use the term preconditioning for physiological conditioning and introduce a new term, behavioral tolerance for diminished avoidance.

Recent studies in *C. elegans*, including ours provided evidence that pathogen and toxin induced stresses simultaneously stimulate cytoprotective responses and aversive behavior (*26–28*). In this study, we set out to investigate how the induction of systemic cytoprotective molecular defenses influence stress-induced aversive behavior and learned behavioral decisions. To this end, we employed two food-derived volatile odors, benzaldehyde (BA) and diacetyl (DA), which are attractive at low, but aversive at high concentrations (*29, 30*). The advantage of these odors is that they contain both the chemosensory cue as well as a dual, attractive or aversive property. Our results indicate a critical role of the ability to mount specific somatic cytoprotective responses in shaping adaptive stress-induced and future behavioral decisions based on associative learning.

## Results

### Concentrated benzaldehyde and diacetyl induce toxicity and food avoidance

Low concentrations of food odors are attractive to *C. elegans*, whereas high concentrations induce an aversive response (*30*). Specifically, worms exhibit a biphasic chemotaxis curve towards concentrated 100% benzaldehyde (ccBA) called benzotaxis (Nuttley et al., 2001 and Fig. S1A). We hypothesized that the second, aversive phase is a defensive behavioral response to ccBA toxicity. Indeed, we found that longer ccBA exposures using the aversive concentration ranges induced extensive paralysis in a dose- and time-dependent manner (Fig. 1A). To investigate whether another concentrated food odor may induce toxicity at aversive concentrations, we tested the chemically unrelated diacetyl. Undiluted diacetyl (ccDA) also triggered biphasic chemotaxis behavior (Fig. S1B) and dose-dependent paralysis at approximately four-fold higher doses compared to ccBA (Fig. 1B). Furthermore, aversive, but lower doses of ccBA and ccDA both impaired thermotolerance (Fig. S1C), demonstrating compromised organismal integrity and stress resistance in response to odor toxicity. Therefore, we used non-paralyzing doses of odors as a source of toxic stress throughout this study.

**Fig. 1.**
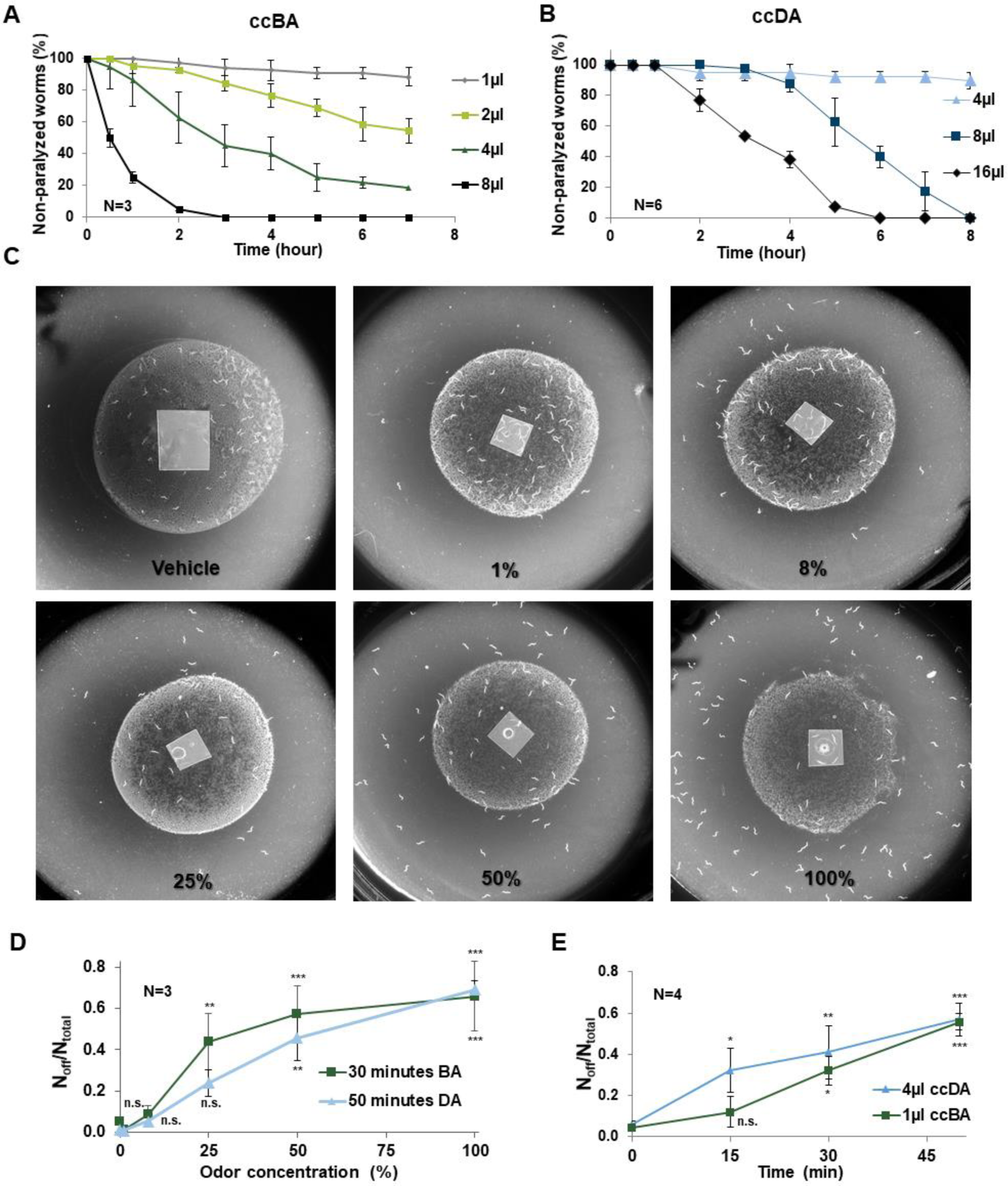
Concentrated benzaldehyde (ccBA) and diacetyl (ccDA) induce toxicity and food avoidance in the aversive concentration range. Time-dependence curves of paralysis to various doses of ccBA (A) or ccDA (B) using a hanging drop assay. (C) Representative images of food leaving behavior in response to various concentrations of BA. BA was placed in ethanol vehicle in a total volume of 1µl in the middle of bacterial lawn. (D) Dose dependence of BA and DA induced food avoidance. BA or DA was placed in a total volume of 1 µl or 4 µl in the middle of bacterial lawn. (E) Time dependence of ccBA and ccDA induced food avoidance. Data are expressed as mean ± SEM, N = number of independent experiments. *p < 0.05; **p < 0.01; ***p < 0.001. Mean durations of odor exposure that induced 50% paralysis by log rank (Mantel-Cox) test were as follows: ccBA – 2 μl: 5.27 h ± 0.17 hours, 4 μl: 2.94 ± 0.21 hours, 8 μl: 0.94 ± 0.14 hours. ccDA? – 8 μl: 5.68 ± 0.20 hours, 16 μl: 3.46 ± 0.17 hours. P-values compared to 1 μl BA or 4 μl DA treatments are <0.001 in all conditions.

Although food is not necessary for adult worms’ survival, worms are continuously feeding and seldom leave the bacterial lawn under laboratory conditions (*31*). Therefore, to establish a more stringent test for behavioral adaptation, we placed a ccBA drop in the middle of an *E. coli* OP50 lawn and monitored food avoidance. We observed that while unexposed worms remained on the lawn after 50 minutes, the majority of ccBA-exposed worms left the food (Fig. 1C). We observed similar, concentration- and time-dependent food aversion phenotypes with both ccBA and ccDA (Fig. 1D, E). These findings indicate that the perception of toxic stress initiates a decision to leave the lawn, giving up the benefit of nutrients for the protection of physical integrity.

### Opposing behavioral patterns elicited by toxic benzaldehyde and diacetyl exposure

We observed that transient ccBA and ccDA exposure increased motility (Fig. S2A), indicating that perception of toxic stress increases locomotor activity which may help instantly avoid the threat. Interestingly, the increased motility returned to baseline after removing ccBA, but showed a sustained elevation after the removal of ccDA (Fig. S2A). Moreover, we found that after an extended, 2-hour exposure to ccBA, animals started to return to the bacterial lawn, whereas the same exposure to ccDA further increased aversion (Fig. S2B). Thus, the adverse physiological effects of ccBA might be eliminated faster than those of ccDA. We reasoned that a preconditioning exposure might differentially affect behavior. To test this, after exposure to the same sublethal doses of odors we investigated the lawn avoidance behavior of naive and preconditioned worms (Fig. 2A). Indeed, we found that preconditioning with ccBA diminished ccBA-induced aversion, while that with ccDA further increased avoidance of ccDA (Fig. 2A, B). For the increased capacity of worms to remain in the presence of toxic ccBA we coined the term “behavioral tolerance”, to the analogy of physiological stress tolerance. Thus, ccBA preconditioning induces behavioral tolerance, while ccDA preconditioning induces sensitization. The effect developed by a 2-hour preconditioning and was only moderately altered by further increasing the duration of the preconditioning exposure (Fig. S2C and D).

**Fig. 2.**
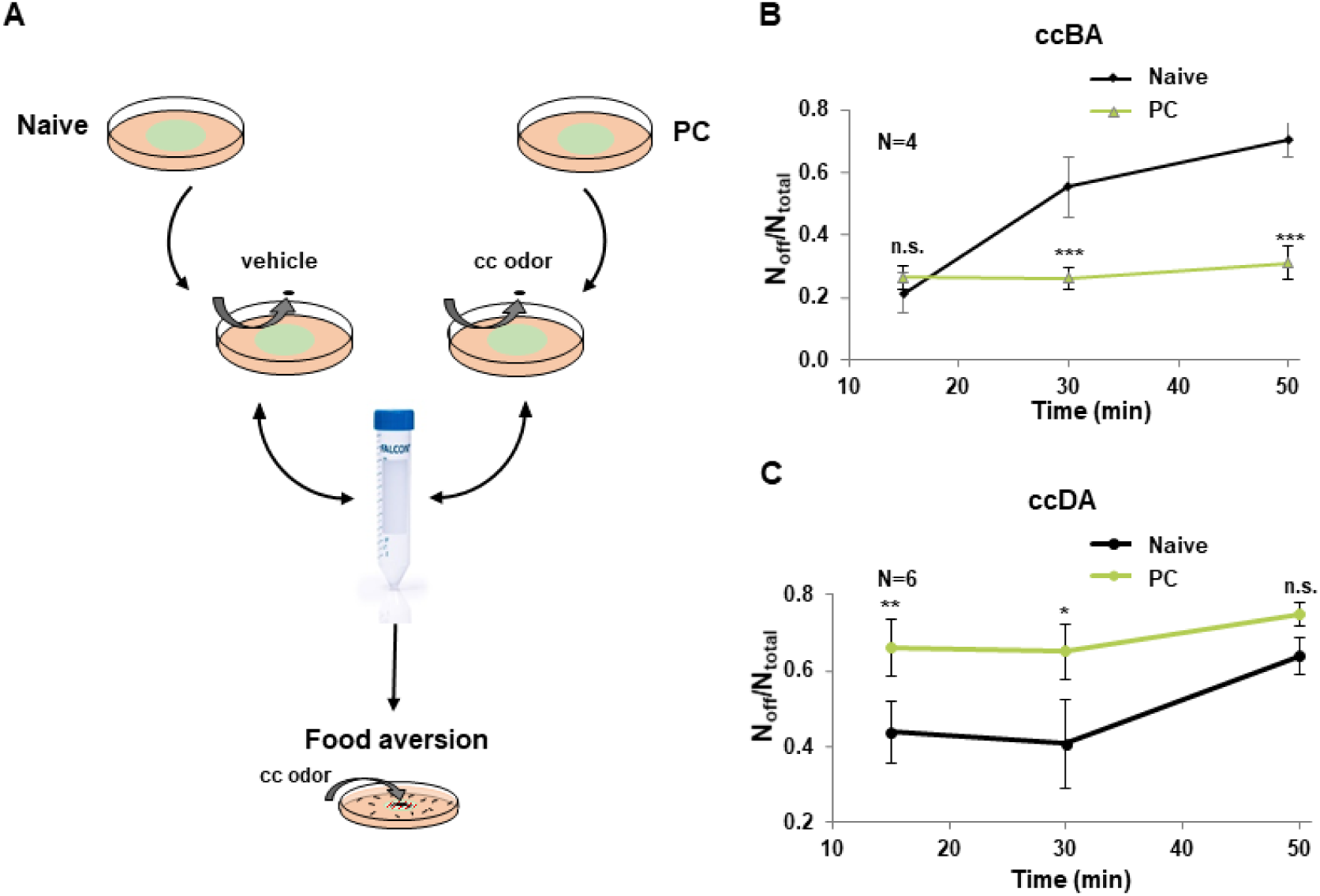
ccBA preconditioning induces behavioral tolerance, while ccDA preconditioning induces sensitization. (A) Experimental setup for preconditioning and food aversion test. Animals were exposed to a hanging drop of concentrated odor (preconditioned, PC) or vehicle (naive), washed and assayed for food aversion. (B) ccBA induced food aversion of naive and ccBA PC animals at different time points. (C) ccDA induced food aversion of naive and ccDA PC animals at different time points. Data are expressed as mean ± SEM, N = number of independent experiments. *p < 0.05; **p < 0.01; ***p < 0.001.

### Concentrated benzaldehyde, but not concentrated diacetyl, activates specific systemic cytoprotective responses

In agreement with our findings on the toxicity of ccBA, previous studies demonstrated that BA induced oxidative stress (*32, 33*). Therefore, we tested various oxidative stress response pathways that might be involved in the organismal adaptation to ccBA. Using the TJ356 strain expressing GFP-tagged DAF-16, we observed that ccBA exposure induced a strong nuclear translocation of DAF-16, comparable to that induced by heat stress. However, DAF-16 remained cytosolic in response to ccDA (Fig. 3A and Fig. S3A). The shift in DAF-16 localization exhibited a clear BA dose-dependence (Fig. S3B). These congruent changes in DAF-16 translocation and food avoidance (cf. Fig. 1D) indicate a potential link between cytoprotective responses and behavioral tolerance.

**Fig. 3.**
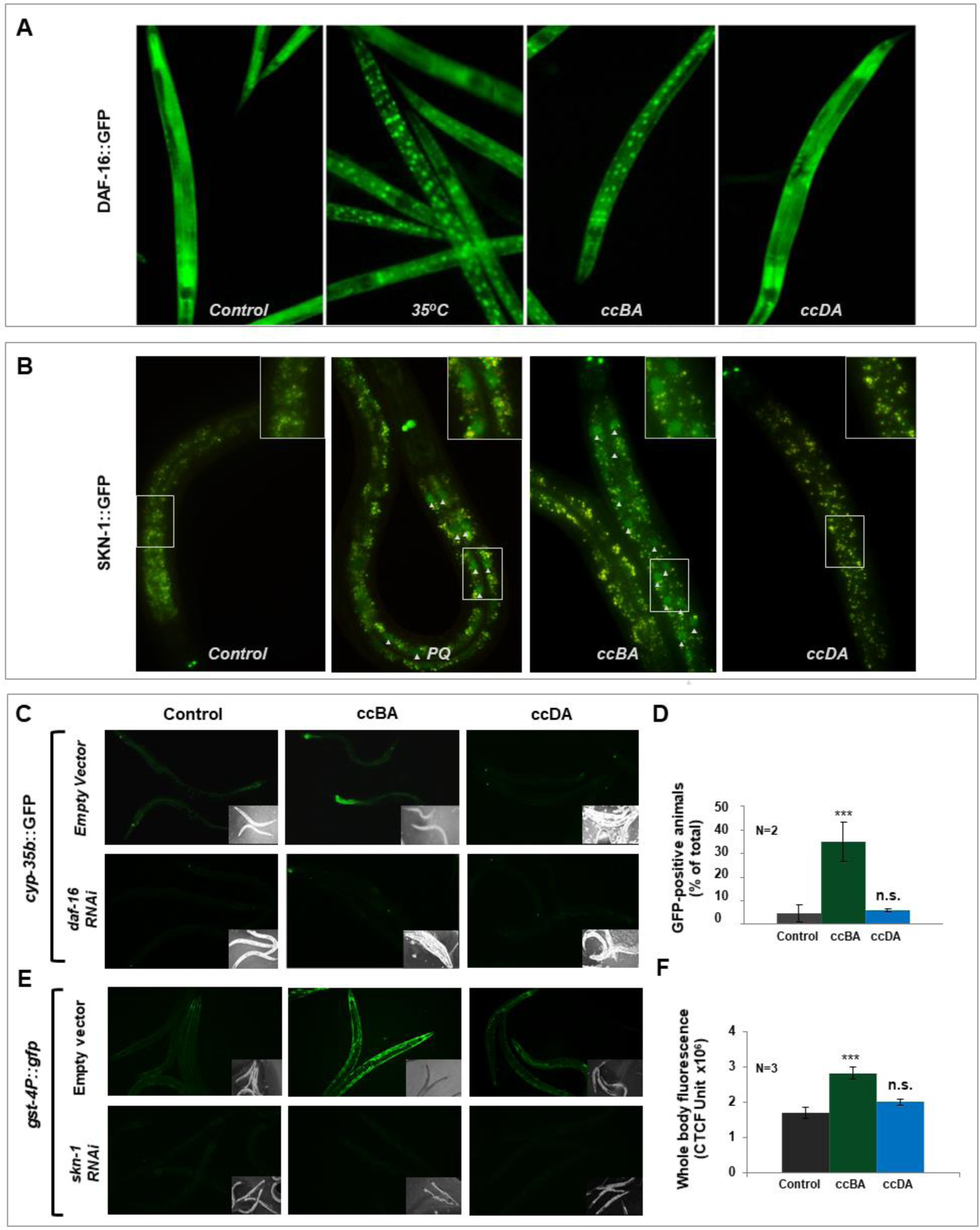
ccBA, but not ccDA, activates specific systemic cytoprotective responses. Representative epifluorescent microscopic images of DAF-16::GFP nuclear translocation in response to heat stress, ccBA or ccDA (A), as well as *SKN-1::GFP* nuclear translocation in response to paraquat, ccBA or ccDA (B, arrowheads). Representative epifluorescent microscopic images (C) and quantification (D) of *cyp-35b::GFP* expression in response to ccBA or ccDA in worms fed by control EV and *daf-16* RNAi. Representative epifluorescent microscopic images (E) and quantification (F) of *gst-4::GFP* expression in response to ccBA or ccDA in nematodes fed by control EV and *skn-1* RNAi. Data are expressed as mean ± SEM, N = number of independent experiment replicates. n.s.: not significant, *p < 0.05; **p < 0.01; ***p < 0.001.

Next, we tested several other stress and detoxification pathways using GFP-tagged marker strains. Translocation of the oxidative-xenobiotic stress master regulator SKN-1::GFP in the LD001 strain was induced by ccBA, but not by ccDA, comparable to that seen upon the oxidative agent paraquat (PQ) treatment (Fig. 3B). Further, ccBA, but not ccDA induced the expression of the phase I marker cytochrome P450 enzyme *cyp-35b* and the phase II enzyme *gst-4* (Fig. 3C-F). The induction of *cyp-35b* was abolished by *daf-16* RNAi, while that of *gst-4* was abolished by *skn-1* RNAi (Fig. 3D, F). Importantly, neither the HSF-1 and DAF-16 target *hsp-16.2*, and the HSF-1 target and endoplasmic reticulum unfolded protein response (UPR) reporter *hsp-4*, nor the SKN-1 dependent *gcs-1* and the DAF-16 dependent *sod-3* reporter was induced by ccBA (Fig. S3C). These findings demonstrate that a specific stress and detoxification response involving a subset of DAF-16 and SKN-1 activated genes is required for the molecular defense against ccBA toxicity.

### ccBA-induced cytoprotective responses confer behavioral tolerance to ccBA, but not to ccDA

We asked whether the cytoprotective responses induced by ccBA which are known to induce physiological stress tolerance might play a role in the generation of behavioral decisions. To this end, we preconditioned N2 and *daf-16* null mutant nematodes with ccBA and studied their food avoidance to ccBA. We found that naive *daf-16* mutants showed avoidant behavior comparable to wildtype, however, they failed to decrease their aversion in response to preconditioning (Fig. 4A). Similar phenotype was obtained by silencing the evolutionarily conserved molecular chaperone Hsp90, which was shown to regulate DAF-16 activity (*34*) (Fig. 4B). Likewise, *skn-1* silencing similarly prevented the development of behavioral tolerance, whereas the activation of SKN-1 by knocking down the WDR-23 protein responsible for its degradation (*35*) augmented behavioral tolerance towards ccBA (Fig. 4C). In sharp contrast, after ccDA preconditioning, neither *skn-1*, nor *wdr-23* RNAi altered the behavioral sensitization towards ccDA exposure (Fig. 4D). RNAi did not silence neuronal Hsp90 and SKN-1 isoforms (Papp et al., 2012; Somogyvári et al., 2018 and data not shown) in agreement with its inability to enter neurons (*36*). These results indicate that specific cytoprotective responses of somatic cells induced by toxic ccBA exposure actively participate in the development of behavioral tolerance.

**Fig. 4.**
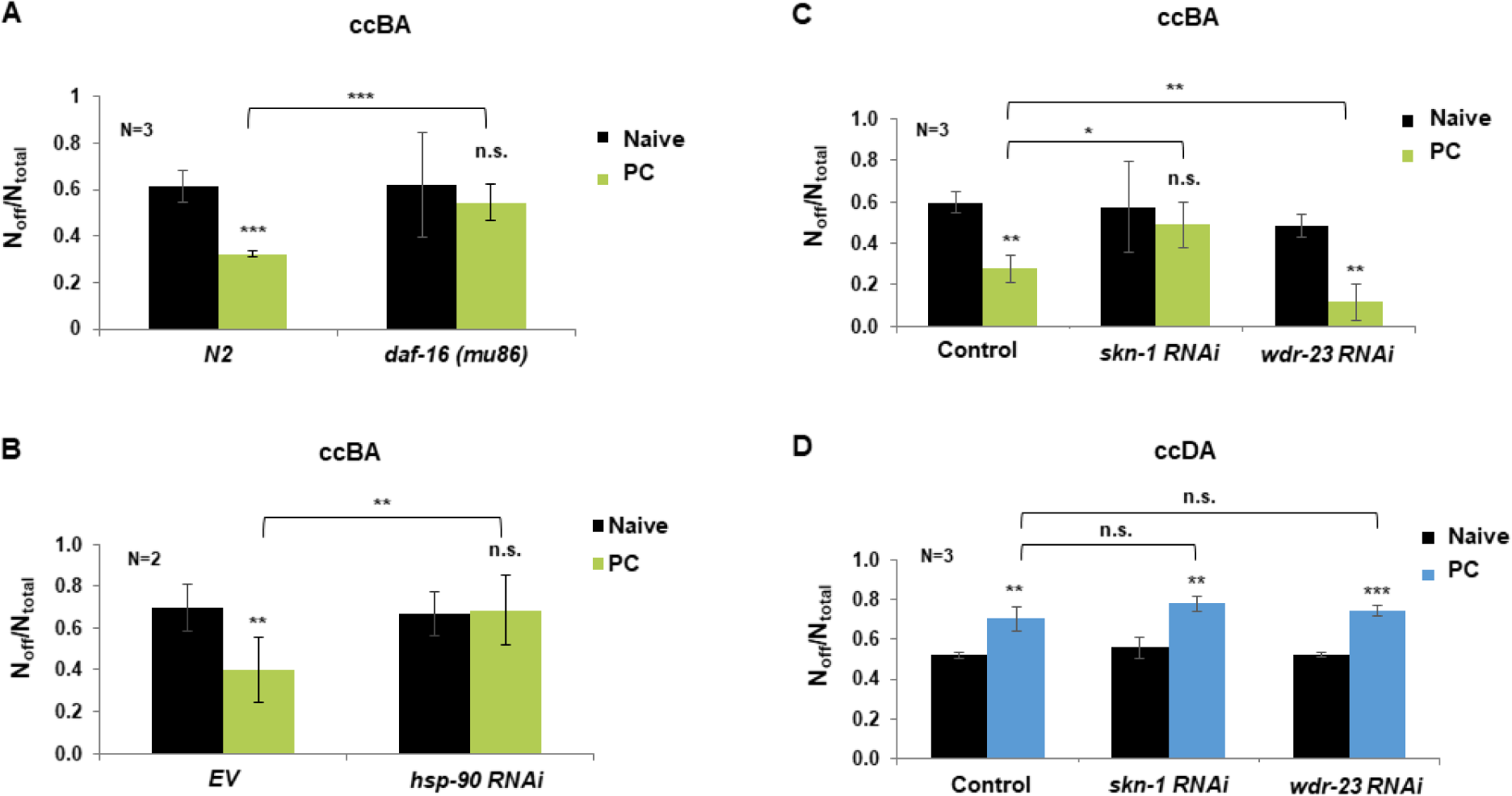
ccBA-induced cytoprotective responses in somatic cells confer behavioral tolerance to ccBA, but not to ccDA. (A) ccBA induced food aversion of naive and ccBA preconditioned N2 wildtype and *daf-16(mu86)* mutant animals, as well as (B) N2 fed by EV and *hsp-90* RNAi bacteria. (C) ccBA induced food aversion of naive and ccBA preconditioned nematodes fed by control EV, *skn-1* and *wdr-23* RNAi, respectively. (D) ccDA induced food aversion of naive and ccDA preconditioned nematodes fed by control EV, *skn-1* and *wdr-23* RNAi, respectively. Data are expressed as mean ± SEM. N = number of independent experiments. *p < 0.05; **p < 0.01; ***p < 0.001.

### Behavioral cross-tolerance is mediated by chemical structure-specific cytoprotective responses

Xenobiotic-induced stress and detoxification responses are related to the chemical structure of the toxin as well as the nature of damage they induce (*37*). We reasoned that the observed BA-dependent cytoprotective machinery might also be induced in response to a chemically similar toxic compound. BA is both spontaneously as well as enzymatically oxidized to benzoic acid during its detoxification (*33, 38, 39*) (Fig. 5A). The chemical structure of the volatile plant stress hormone methyl-salicylate (MS) (*40*), harboring an aromatic benzene ring and an esterified carboxyl group is closely related to that of benzaldehyde and benzoic acid (Fig. 5A). We found that similarly to ccBA, concentrated MS (ccMS) was also toxic and induced food avoidance behavior (Fig. S4A and B). Moreover, ccMS and ccBA shared identical molecular defense responses, including DAF-16 translocation, *cyp-35b::GFP* and *gst-4P::gfp* expression (Fig. 5B). Importantly, preconditioning with either ccMS or ccBA reduced food aversion in response to a subsequent ccMS exposure (Fig. 5C). However, ccBA preconditioning did not affect food aversion in the presence of ccDA, indicating that DAF-16 and SKN-1 dependent processes are unable to reduce ccDA toxicity (Fig. 5D). We conclude that the BA-specific cytoprotective responses confer behavioral cross-tolerance towards a toxin harboring a similar chemical structure, but not towards another compound, DA, which is unrelated chemically and probably by mechanism of action.

**Fig. 5.**
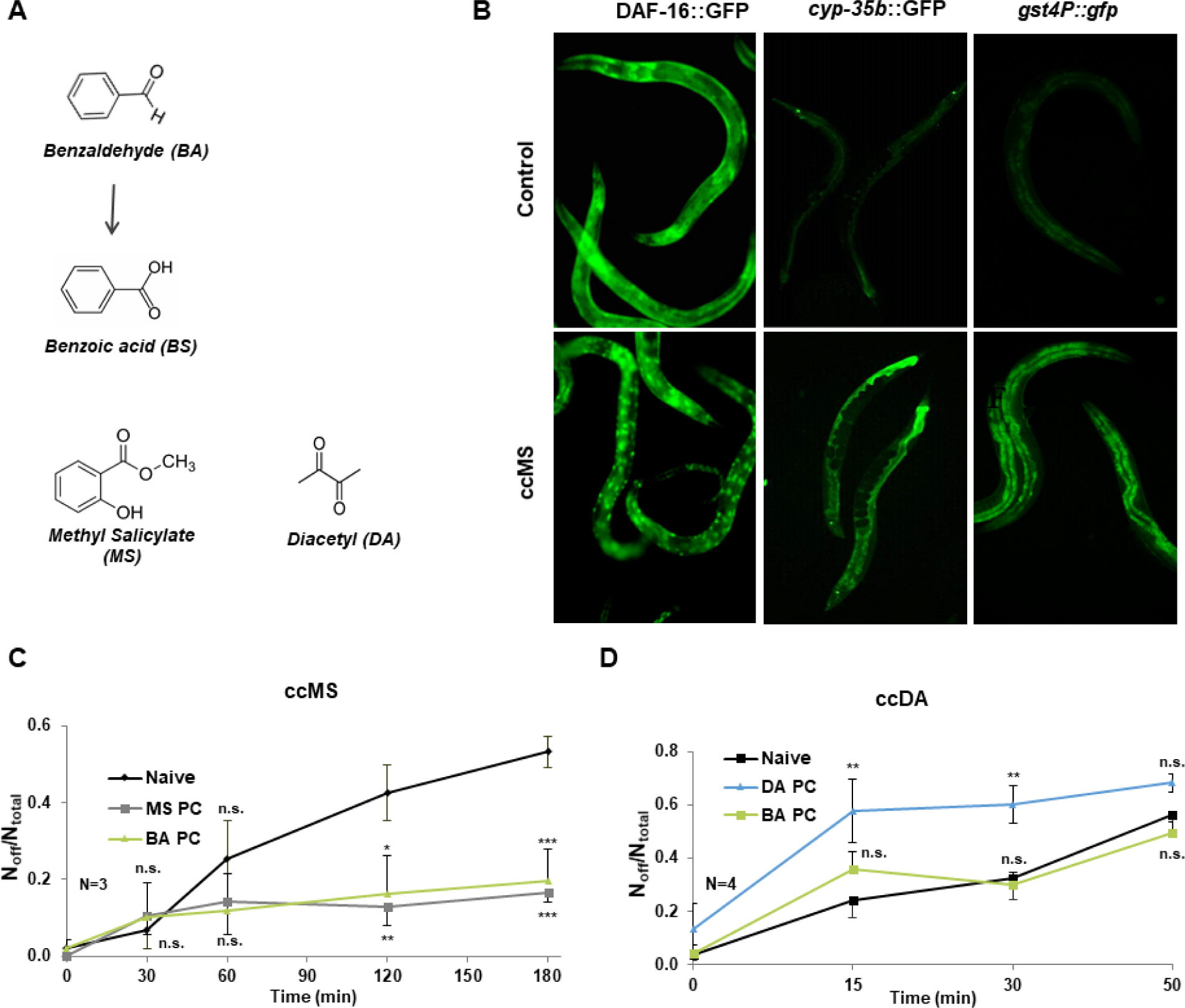
Behavioral cross-tolerance is mediated by chemical structure-specific cytoprotective responses. (A) Benzaldehyde, its metabolite benzoic acid and methyl-salicylate (MS) share similar chemical structures. (B) Representative epifluorescent microscopic images showing the effect of concentrated MS (ccMS) exposure on DAF-16::GFP nuclear translocation, *cyp-35b*::GFP and *gst-4P::gfp* reporters expression. (C) ccMS (MS PC) as well as ccBA (BA PC) preconditioning significantly and comparably decrease ccMS-induced lawn avoidance, while ccBA preconditioning does not reduce ccDA-induced lawn avoidance (D). Error bars represent mean ± SEM compared to the respective naive values, N = number of independent experiments. n.s.: not significant, **p<0.01, ***p<0.001.

### Deficient or efficient cellular defenses generate relevant learned behaviors to stress-associated olfactory cues

The lack of behavioral tolerance in case of ccDA preconditioning indicates inefficient cellular protection, in agreement with our findings (see Fig. 3). Moreover, the phenomenon of behavioral sensitization, a significantly faster and more pronounced aversive response towards the odor suggests a role for avoidant associative learning. To test this, we investigated alterations in behaviors towards attractive (1%) doses of DA and BA after pre-exposure of toxic, concentrated doses of the respective odors (Fig. 6A). Indeed, worms preconditioned with ccDA significantly reduced their chemotaxis towards naturally attractive 1% DA (Fig. 6B) and even chose to leave the food in the presence of 1% DA (Fig. 6C). We also investigated decision making by providing both DA and BA naturally associated with food in an odor choice assay. The aversive change of the DA olfactory cue was underscored by an almost complete shift in odor preference to BA (Fig. 6D). In contrast, worms preconditioned with ccBA did not leave the bacterial lawn in the presence of 1% BA (Fig. 6F). Moreover, they maintained their chemotaxis towards, 1% BA (Fig. 6E), when the olfactory cue of BA was the only option. However, they displayed reduced preference to BA in the simultaneous presence of attractive DA (Fig. 6G). These results are consistent with the formation of distinct, avoidant or tolerant learned behaviors associated to the sensory cues of DA and BA, respectively, after a previous encounter with their toxic doses, which appear to stem from the prior internal experience resulting from a deficient or efficient cytoprotection.

**Fig. 6.**
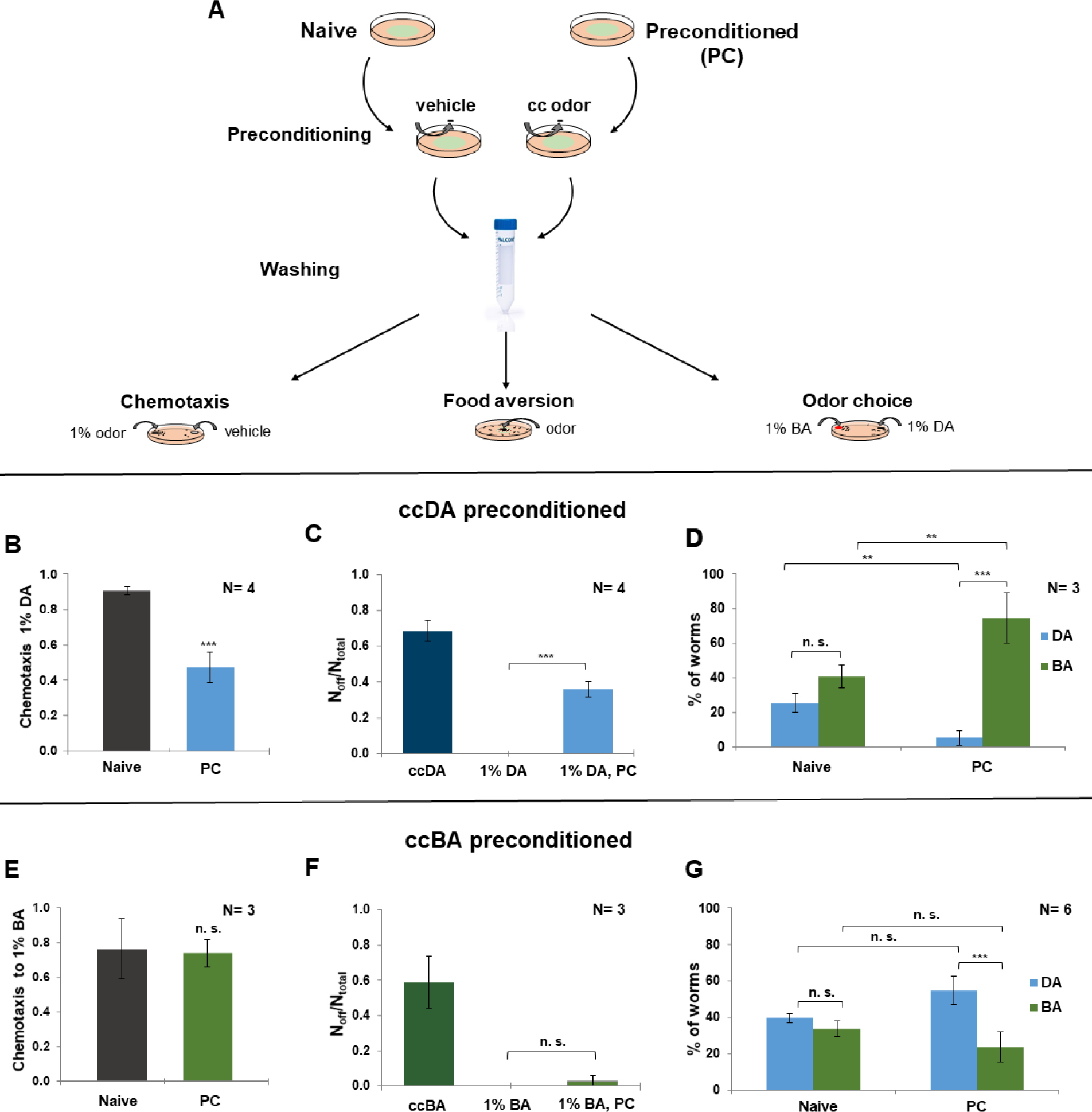
Learned stress-associated behaviors are shaped by prior cellular defenses against stress. (A) Experimental design for toxic odor preconditioning induced associative learning. Animals were exposed to a hanging drop of concentrated odor (ccBA or ccDA, preconditioned, PC) or vehicle (naive), washed and assayed for chemotaxis, food aversion and odor preference using 1% of the odors. ccDA preconditioning significantly reduces chemotaxis to (B), and stimulates lawn avoidance from (C) 1% DA, and entirely shifts odor preference to 1% BA (D). ccBA preconditioning does not affect chemotaxis to (E), and lawn avoidance from (F) 1% BA, but induces a modest reduction of preference to 1% BA in the presence of 1% DA (G). Odor choice was quantified by scoring worms on BA and DA spot. Error bars represent mean ± SEM. N = number of independent experiments. n.s.: not significant, *p < 0.05; **p < 0.01; ***p < 0.001.

### A stress-associated memory of somatic resilience enables context-dependent decision making

The association of naturally attractive odors with learned stress-reactive behaviors raises the possibility that the learned experiences may give rise to associative memories to cope with a similar anticipated future insult. On the other hand, forgetting irrelevant experiences is also important as both the organism and the environment is changing. Indeed, a 2-hour recovery period after a single ccBA preconditioning of two or four hours significantly attenuates behavioral tolerance in the food leaving assay (Fig. 7B). We reasoned that repeated exposure of the same conditions might reinforce the co-occurring experience resulting in a lasting neural representation. To test this, we employed a protocol of massed training known to induce short and intermediate term associative memories (STAM and ITAM) (*41, 42*) (Fig. 7A). By definition, STAM decays within, whereas ITAM persists over, one hour. Massed training of one-hour exposure to ccBA four times resulted in a potent behavioral tolerance that was retained after a 2-hr recovery (Fig. 7C). A single 4-hour preconditioning, a physiological stress of the same duration induced comparable food aversion immediately after training (cf. Figs. 6F and 7C), which indicates similar levels of cytoprotection and makes it unlikely that the behavioral tolerance is a result of higher physiological stress tolerance. Likewise, ccDA massed training induced a robust food avoidance in the presence of 1% DA which persisted after 2-hr recovery (Fig. 7D). Again, massed-trained worms exhibited comparable aversive behaviors against 1% DA as if they encountered ccDA (cf. Fig.s 2C, 6C and 7D). These results are consistent with the reinforcement of toxic stress-associated neurosensory integration into different associative memories of active coping or passive avoidance. The stability of memory after two hours indicates the formation of ITAM. Finally, we asked how the coping memories affect the choice between the stress-associated and a natural attractive odor olfactory cue. Massed training with ccBA almost entirely shifted the preference towards DA (Fig. 7E), potentiating the change already observed by a single preconditioning (Fig. 6G). This phenomenon also shows an apparent similarity to the complete disappearance of DA preference after a single preconditioning with ccDA (cf. Fig.s 7E and 6D). Nonetheless, in contrast to the compelling avoidant behavior to the memory of uncompensated harm, the memory of cellular protection not only provides the ability to cope with anticipated toxicity for food, but allows a context-dependent decision to spare resources when the organism also perceives the cue of a potentially toxin-free food. This result also suggests that the memory of a stressful insult contains the representation of the original valence of the olfactory cue, the internal experience of stress-induced harm and that of the activated physiological protection.

**Fig. 7.**
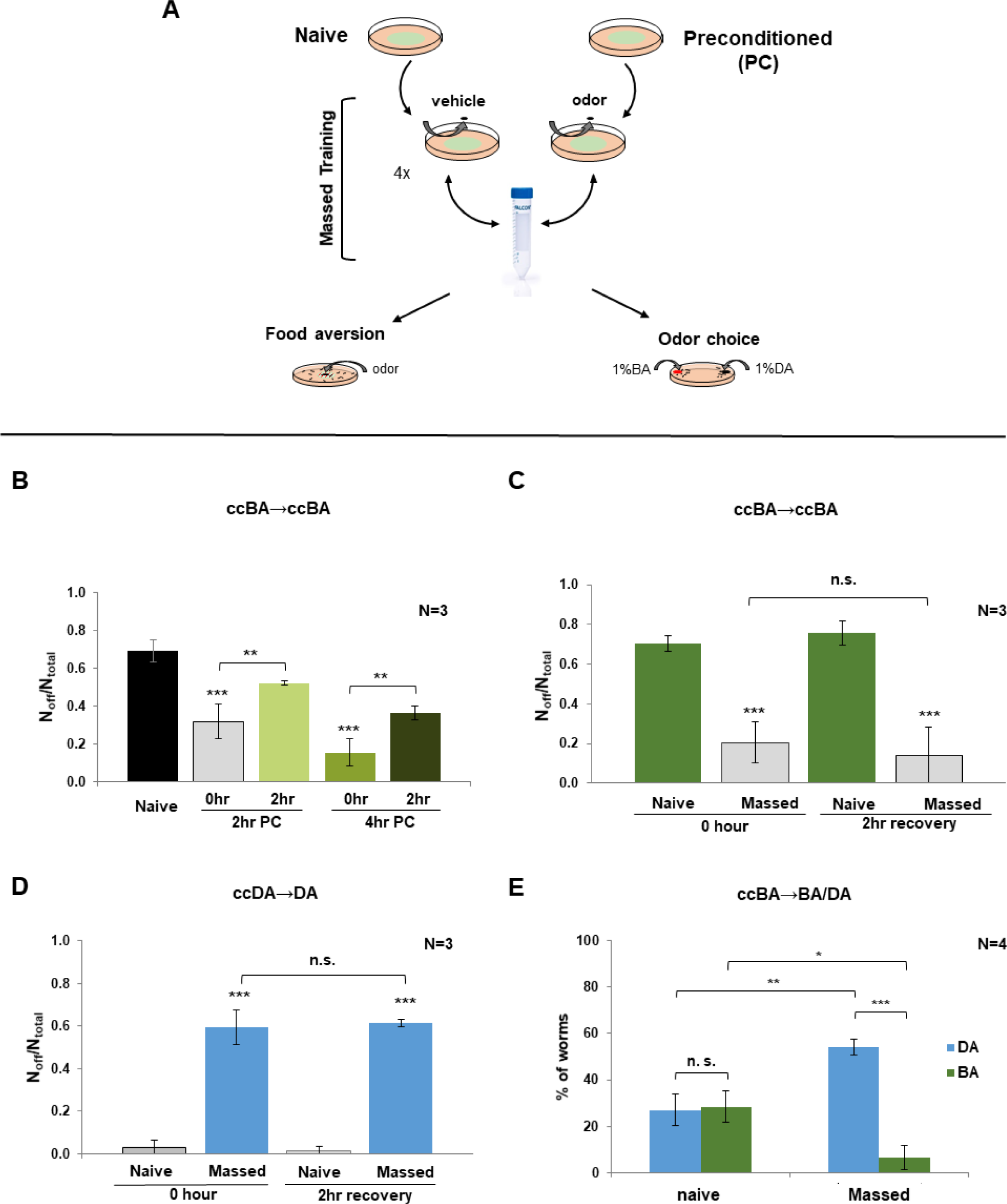
Reinforcement of stress-associated internal experiences form distinctive intermediate term associative memories. (A) Experimental design for toxic odor preconditioning induced associative memory. Animals were exposed to a hanging drop of concentrated odor (1 µl ccBA or 4 µl ccDA, preconditioned, PC) or vehicle (naive), using a single preconditioning (B) or a 4x massed training protocol (C-E), then assayed for food aversion or odor preference immediately or after the indicated recovery periods. (B) A 2-hour recovery period decreases the food aversion elicited by a single 2-hour or 4-hour ccBA preconditioning. (C) Nematodes exposed to a 4×1-hour ccBA massed training retain the behavioral tolerance to ccBA after a 2-hour recovery period. (D) Nematodes exposed to a 4×1-hour ccDA massed training maintain their avoidant behavior to 1% DA after a 2-hour recovery period. (E) A 4×1-hour ccBA massed training induces a robust shift in odor preference towards 1% DA. Odor choice was quantified by scoring worms on BA, DA and on the empty agar surface (0). Error bars represent mean ± SEM, N = number of independent experiments. n.s.: not significant, *p < 0.05, **p<0.01, ***p<0.001.

## Discussion

In this study we have set up a paradigm in *C. elegans* to assess the impact of cytoprotective responses on behavioral decisions. We have shown that the innately attractive odors benzaldehyde and diacetyl, when employed at high concentrations, induce toxicity and behavioral aversion of the odor-contaminated bacterial lawn. ccBA-induced somatic cytoprotective responses involving DAF-16, SKN-1 and Hsp90 conferred behavioral tolerance to ccBA and cross-tolerance to concentrated methyl-salicylate (ccMS), while neither behavioral tolerance nor apparent molecular defenses were observed upon exposure to ccDA. Massed training generated an associative memory that made diluted DA aversive but enabled animals to decide whether to approach or to avoid diluted BA depending on alternative choice. Our study suggests that the (in)ability of *C. elegans’* somatic cells to counteract toxic stress with cytoprotective mechanisms regulates behavior during stress and determines learned behavioral decisions upon re-encounter with stress-associated olfactory cues (Fig. 8).

**Fig. 8.**
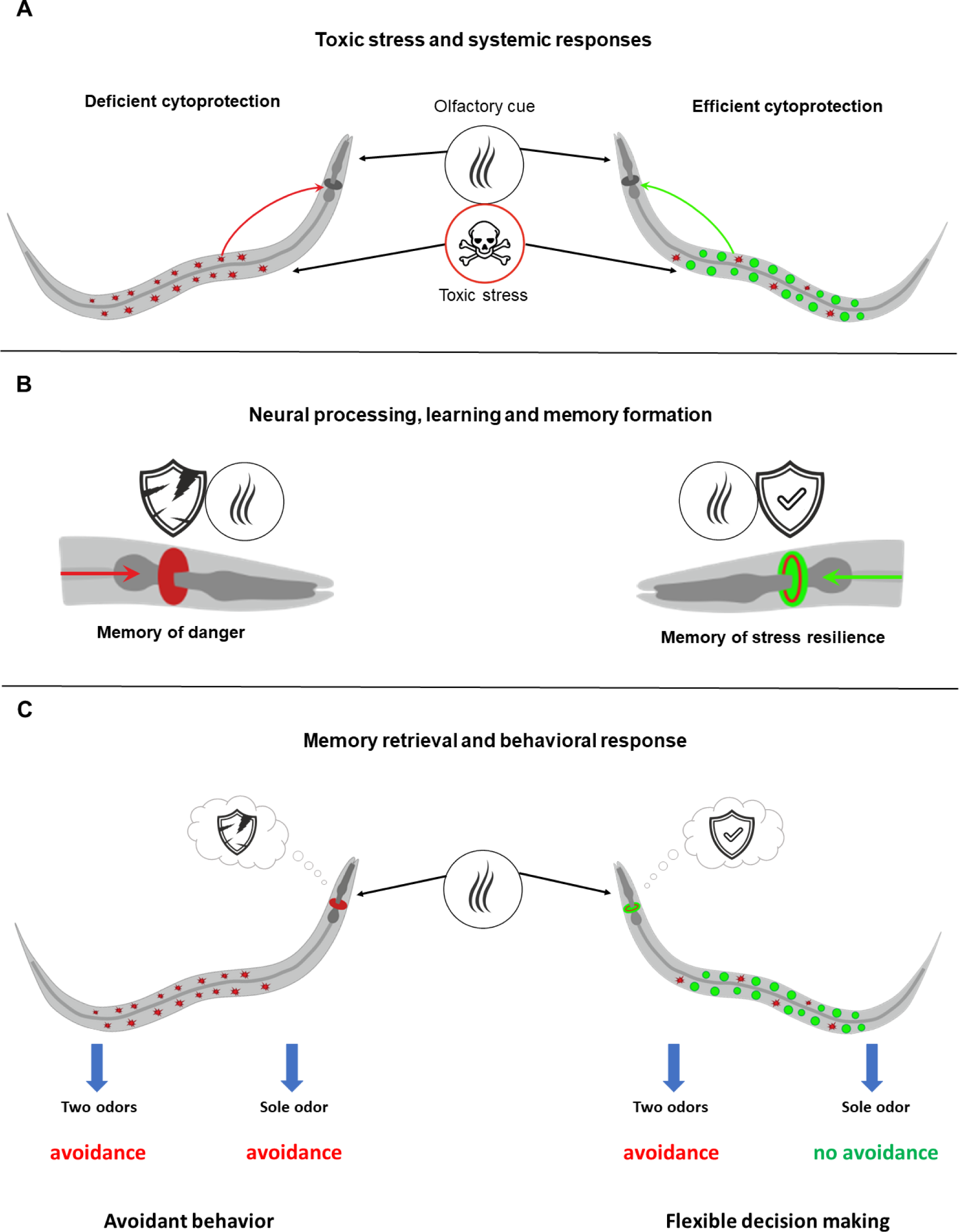
Model for the regulation of learned behavioral decisions by somatic cytoprotective responses. (A) Toxic stress-induced disturbance of cellular homeostasis (red symbols) prevails in the absence of adequate cytoprotection (left) and emits danger signals (red arrow) towards the nervous system (dark grey nerve ring in the head). Stress-specific cytoprotective responses (right) restore cellular homeostasis (green symbols) and suppress danger signals (green arrow). (B) Simultaneously, signal processing in the nervous system generates an internal experience of danger (damaged shield) or stress-and-protection (intact shield). Integration with the co-occurring olfactory cues forms either a danger-based (red nerve ring) or stress-resilient (red-green nerve ring) associative memory. (C) Memory retrieval by the respective olfactory cues evokes either a stereotypical avoidant behavior or a flexible behavioral decision depending on the external context, such as the absence or presence of another food-indicative odor besides the stress-associated olfactory cue.

Previous studies reported either decreased attraction or aversion to high concentrations of food-derived odors (*15, 29, 30, 43*). One study proposed that the change in odor preference towards concentrated benzaldehyde was due to olfactory adaptation (*29*). Although this mechanism might hold true, several findings of our study argue for an alternative explanation. The *(i)* strong dose-dependent food aversion (Fig. 1C-E) *(ii)* increased motility (Fig. S2A) *(iii)* progressive dose-dependent paralysis (Fig. 1A, B) *(iv)* compromised thermotolerance (Fig. S1C) *(v)* induction of robust systemic cytoprotective responses by ccBA (Fig. 3) *(vi)* the comparable ccBA concentration dependence of aversion and DAF-16 nuclear translocation (Fig. 1D and Fig. S3B) (*vii*) the manipulation of these responses modulates the development of behavioral tolerance to ccBA (Fig. 4) (*viii*) RNAi is unable to penetrate neurons (*36*) all indicate somatic toxicity as an underlying mechanism. This interpretation gains support from the facts that low concentrations of benzaldehyde and isoamyl alcohol mediate attraction via activating the AWC chemosensory neuron, whilst high concentrations activate the polymodal nociceptive ASH neuron, which in turn drives repulsion (*15, 30, 43, 44*). Consistent with our study, various studies in mammals describe the toxic effects of benzaldehyde (*33, 45, 46*) and diacetyl (*47–49*) such as inhalation toxicity and long-term impairment in lung function. Together with the evidence presented by the above reports, our study suggests that tissue damage caused by odor toxicity stimulates aversion (Fig. 8). Perhaps neural (*29*) and somatic (our study) inputs are integrated to increase the robustness of behavior. Our findings also draw attention to nematode associative learning experiments where different conditions are paired with concentrated odors (*42, 50, 51*), because odor toxicity might stimulate repulsion independently of, or synergistically with, the unconditioned stimulus. Therefore, behavioral experiments using diluted odors as the conditioned stimulus are recommended.

Our observations on the toxic odor-induced food aversion (Fig. 1) indicates this neurobehavioral response is a first line of defense against dangerous insults, which preserves physical integrity and spares resources. ccBA exposed worms, however, started to return to food during the second hour and a preconditioning exposure also diminished ccBA avoidance (Fig. S2B and Fig. 2B). Reduced avoidance coincided with DAF-16 and SKN-1 activation and induction of phase 1 and 2 xenobiotic detoxification reporters (Fig. 3), consistent with the aromatic structure and toxic profile of ccBA. Such cytoprotective stress and detoxification responses co-operate to ensure survival, stress tolerance, immunity and longevity (*4, 18, 52, 53*), forming a cellular defense. Our findings that *daf-16* knockout, *hsp-90*, *skn-1* and *wdr-23* RNAi in somatic cells specifically modulate ccBA avoidance (Fig. 4) show that specific stress-responsive regulators control aversion, revealing a novel regulatory role of somatic cytoprotective responses on behavioral decisions.

We found ccBA-induced behavioral cross-tolerance to concentrated methyl-salicylate (ccMS) (Fig. 5C). Our results on identical cytoprotective responses shared by ccBA and ccMS (Fig. 5) suggest that the responses stimulated by ccBA preconditioning also eliminate the toxic agent and repair damage during ccMS exposure. Indeed, high doses of methyl-salicylate cause heavy toxicity in mammals (*54*). Thus, the preservation/restoration of tissue integrity by toxin-specific cytoprotective responses suppresses aversion. Consistent with this idea, ccDA treatment resulted in sustained hypermotility after ccDA removal and increased ccDA aversion (Fig. S2A and Fig. 2C). Furthermore, ccDA did not appear to activate considerable molecular defenses (Fig. 3) and neither SKN-1 manipulations (Fig. 4D) nor the systematic induction of cellular defenses by ccBA preconditioning (Fig. 5D) affected ccDA aversion. These results also exclude the unlikely possibility that induction of systemic cytoprotective responses *per* se inhibits aversive behavior. Hence, in the absence of adequate molecular defenses, the disturbance of cellular homeostasis may represent a danger signal which induces aversion.

The mechanisms by which toxin-induced disturbances in cellular homeostasis elicits behavioral aversion are yet unclear, but the modulation of aversion by somatic RNAi manipulations (Fig. 4) indicates an endocrice response. The stress-activated JNK and p38 MAP kinases are conserved signal transducers of cell stress including xenobiotic, oxidative, proteo-, genotoxic and pathogen stresses (*55*). Indeed, both the JNK-1 ortholog KGB-1 (*56*) and the p38 ortholog PMK-1 (*57, 58*) were shown to monitor homeostasis and transmit signals to induce aversion in response to toxicity and infection. Further, yet unidentified signals emerge from the bloated intestine in response to infection (*27*), the major inner barrier and site of immunity and detoxification. Both DAF-16 and SKN-1 positive nuclei appear to be located around the gut lumen in our experiments (Fig. 3). The apparent impermeability of the cuticle to chemicals (*59*) and the predominant localization of SKN-1 and DAF-16 isoforms in the intestine suggest that signals evoked by odor toxicity are also likely to emanate primarily from the intestine. Neuronal events might involve the NPR neuropeptide receptor and serotonin signaling pathways, which were shown to be required for aversive behavior and aversive olfactory learning against pathogens, toxins and for methyl-salicylate (*27, 56, 60–64*). However, other, yet, unidentified pathways might also be involved in the elicitation of the behavioral responses. We note that worms exposed to ccBA stayed at the edge of the food lawn but deserted at higher concentrations (see e.g. the 8% ccBA and above in Fig. 1C), similarly to the observations made using the repellent 2-nonanone (*65*), suggesting the multisensory integration of attractive and aversive impulses for decision making. Taken together, the elucidation of the mechanisms of cell stress-elicited behavioral decisions is an intriguing subject of future research.

Our findings that after a single stress exposure, characteristic stress-related behavioral responses are retrieved by the associated olfactory cues (Fig. 6) are indicative of associative learning. Similar phenomena, where somatic/cellular stress regulates learned aversion are the compromise in vital functions by toxins or RNAi targeting essential cellular processes and the intestinal bloating caused by pathogen bacteria (*56, 66*). Although the prevalence of new behaviors decreases with time, repeated stress exposures not only maintain (food leaving on diluted DA after ccDA massed training) but also enhance (odor choice after ccBA massed training) the respective behavioral choices (Fig. 7D, E). These findings are consistent with forgetting and intermediate term associative memory formation by reinforcement of the experience (*42, 67*). The elicitation of opposing behaviors by DA and BA olfactory cues after preconditioning demonstrates that the absence or presence of adequate cytoprotective responses at the time of stresses is a critical regulator of future behavioral decisions to anticipated stress (Fig. 8). The internal experience of disrupted tissue homeostasis by ccDA in the absence of cytoprotection is integrated into an associative memory of danger, which reverses the naturally attractive valence of DA and upon retrieval gives rise to avoidant behavior. Efficient cytoprotection in response to ccBA restores homeostasis and forms a memory of protection, which upon retrieval elicits a tolerant behavior. Besides, the increased preference of DA over BA after ccBA preconditioning (Fig. 6G and 7E) suggests that the natural valence of BA is integrated with the cost to maintain it, probably through the associated experience of toxic stress. Such representation allows individuals to consider whether investment of resources for self-protection are needed or not to obtain food. Thus, although the BA-associated memory of somatic resilience creates a flexible choice depending on the local context of alternative food odor, whereas the DA-associated memory of danger gives rise to a stereotypic aversion, both behavioral decisions depend on the respective neural contexts of prior experiences (Fig. 8).

An evolutionarily conserved adaptive response to stress is the “fight-or-flight” response, originally coined for the neuroendocrine system (*3, 68*). Together with recent studies we here show the co-occurrence of behavioral “flight” responses with molecular stress and immune “fight” responses combating stress (*27, 28, 56*). Moreover, our studies reveal a regulatory link between intracellular cytoprotective responses and behavior suggesting a coordinated action of the fight and the flight responses to preserve organismal integrity. Beyond ensuring physical survival, the maintenance of cellular homeostasis by stress responses equips nematodes with behavioral tolerance to stay in or to approach real and anticipated stressful locations. Further, temporary avoidance in stresses that overwhelm molecular defenses allows the restoration of physiological integrity and the strengthening of cytoprotective mechanisms. Moreover, genetically weakened or absent molecular defenses narrow fitness by reinforcing avoidant behavior. Our work implies that memories of past stresses accompanied by insufficient cellular defenses may condition to avoidance. Avoidant behaviors are characteristic to various human mental disorders, such as phobias, panic attacks, complex posttraumatic stress disorder and eating disorders. These diseases are accompanied by intense physical sensations of stress and overwhelming fear and emotions to jettison or to avoid perceived danger, which happen in response to specific or unidentified sensory cues (*11, 69*). The foundations of stress and detoxification responses and learning are conserved between nematodes and humans. Thus, it might be conceivable that unconscious memories of prior stressful somatic experiences govern emotions and behaviors in response to sensory cues.

## Conclusions

This study shows how organisms ensure optimal self-protection during environmental stress by coordinating physiological and neurobehavioral defenses. Specifically, our findings reveal a critical role of somatic defenses in regulating behavioral avoidance and associative learning *via* the activation of conserved cellular stress responses. The mechanism depicted here enables animals to anticipate adverse conditions by retrieving stress memories and tailor their behavioral decisions depending on their past physiological response to the stressor. Whether such cellular memories might shape human behavior is subject of future studies.

## Methods

### Materials

The reagents benzaldehyde, diacetyl and methyl-salicylate were obtained from Sigma Aldrich. ccBA and ccDA abbreviate undiluted (concentrated) benzaldehyde and diacetyl, respectively. All other chemicals were obtained from Sigma or Fluka, if not otherwise mentioned.

### *C. elegans* strains and maintenance

All strains used were provided by the Caenorhabditis Genetics Center: N2 (Bristol) wild type; TJ356 [daf-16p::daf-16a/b::GFP + rol-6(su1006)]; LD001 {Is007 [skn-1::gfp]};TJ375 [hsp-16.2p::GFP]; CF1038 [daf-16(mgDf50)]; CY573 [bvls5(cyp-35B1p::GFP + gcy-7p::GFP)]; MJCU017 {kIs17[gst-4::gfp, pDP#MM016B]X}; LD1171 {Is003 [gcs-1::gfp]}; SJ4005 [hsp-4::GFP]; CF1553 {muIs84[pAD76(sod-3::GFP)]};. Strains were grown and maintained as previously described (*70*). Animals were synchronized by allowing adults to lay eggs for 4 hours. All experiments were performed using day 1 adults, if not otherwise indicated.

### Odor preconditioning and massed training

Preconditioning treatments were performed using the hanging drop method to prevent direct contact of concentrated volatiles with worms in the presence of bacterial food source to prevent the associated experience of starvation. More precisely, 1µl and 4µl drop of concentrated benzaldehyde (ccBA) or diacetyl (ccDA), respectively, was placed on the lid of 6 cm NGM plates seeded with OP50, containing a synchronous population of 200-300 young adults. The plate was sealed with parafilm to maintain a relatively constant dose of volatile. If not otherwise stated, preconditioning time was three hours. Massed training protocol was designed as described (*41, 42*) employing four sequential one-hour exposures to hanging drops of 2 µl ccBA, 4µl ccDA or vehicle with intermittent washes in M9 buffer.

### Acute toxicity and thermotolerance measurements

Toxicity and thermotolerance assays were carried out at 20°C, and at 35°C, respectively, by using approximately 25-40 worms per plate in three replicates in 3cm NGM plates in case of toxicity, and 6cm NGM plates in case of thermotolerance. Both toxicity and thermotolerance were measured by counting paralyzed worms using „head lifting” behavior of moveless animals (*71*). If an apparently paralyzed worm was not able to display at least “head lifting” movements following gentle fall of assay plate into experimental surface, it was counted as „paralyzed”. Paralysis index was calculated as the average of N_paralyzed_/N_total_ at each time point. Animals that crawled off the agar surface were censored.

### Chemotaxis assays

Chemotaxis experiments were carried out as previously described (*15*) and carried out earlier by our lab (*72*), with modifications. Briefly, synchronous population of young adults were washed twice in M9 buffer, then 80-100 worms were placed in the middle of a 10 cm CTX assay plate containing the odors without anesthetics in order to monitor the actual decisions at indicated time points. In kinetic chemotaxis, plates were streaked at each centimetre to measure the weighted distribution of worms at indicated time points. The Weighted Chemotaxis Index (WCI) were calculated as previously described (*29*).

### Food avoidance assay

Bacterial lawn-avoidance experiments were performed as previously conceived (*73*), with modifications. Briefly, 50-80 synchronous day-1 adults were washed twice with M9 buffer and dropped onto the OP50 lawn in the middle of 6 cm NGM plates. Worms were allowed to settle for 30 minutes, unless otherwise indicated. A drop of given odor were placed on a piece of parafilm in the middle of the OP50 lawn. Animals on or off the lawn were counted at each indicated time point. Worms incapable to move or crawled off the agar surface were censored. Food-leaving index was calculated as the average of N_off_ /N_total_ taken from three technical replicates.

### Motility assay

Motility was characterized as described (*74*) and performed earlier (*72*) by counting body bends for 1 minute using 10-15 animals in each condition. After measuring baseline motility on an OP50-seeded NGM agar plate, a toxic dose of odor hanging drop was placed on the lid and motility was measured at the indicated time points.

### RNA interference

RNAi strains were obtained from Source Bioscience (Notthingam, UK). RNAi treatments were performed as previously described (*75*). RNAi feeding clones were grown overnight in LB medium containing 100 μg/ml ampicillin. Worms were grown on plates containing 1 mM IPTG, 50 μg/ml ampicillin and 6.25 μg/ml tetracyclin and seeded with *E*. *coli* HT115 strains harboring the L4440 empty vector (EV) control and specific RNAi vectors, respectively, from hatching.

### Fluorescence microscopy

Analysis and quantification of fluorescence was carried out as previously described (*24*), with modifications. After treatments, at least 20 worms per condition were picked individually and immobilized by 20 mM NaN_3_ washed in M9 buffer onto a 2% agarose pad. Microscopic examination was carried out on a NIKON Eclipse E400 type fluorescence microscope linked to a Diagnostic Instruments SPOT 500 camera in case of TJ356, TJ375, CY573, MJCU017, LD1171, SJ4005, CF1553 strains; and OLYMPUS CKX53 Fluorescence microscope, OLYMPUS DP74 Cooled color camera in case of LD001 strain, using green fluorescent filters. Images are representatives of at least three independent experiments. Fluorescence intensity measurements were quantified with ImageJ. Visualization of skn-1::GFP nuclear punctae were carried out by OLYMPUS CellSens v2.3 Imaging software.

### Odor preference assay

Odor preference was carried out in standard CTX plates. 80-100 naive and preconditioned young adults were washed off twice in M9 buffer and dropped into the middle of the assay plate. Odors were placed into two sides of the assay plate and worms were allowed to migrate for 50 minutes. Data are expressed as % of animals migrated into a 1 cm drawn circle around the respective odors as well as % of animals remained out of circles.

### Statistical analysis

Kaplan–Meier log-rank tests using the program IBM SPSS Statistics were carried out to evaluate toxicity assays. Food avoidance and chemotaxis assays were examined by one-way ANOVA with Fisher’s LSD post-hoc test. Odor preference assays were analyzed by two-way ANOVA with Fisher’s LSD post-hoc test after evaluation of normal distribution significance by Shapiro-Wilk test. Significance in fluorescence intensity were calculated by unpaired Student’s t-test following evaluation of normal distribution significance by Kolmogorov-Smirnov test and Shapiro-Wilk test. One-way ANOVA with Fisher’s LSD post-hoc tests, Shapiro-Wilk and Kolmogorov-Smirnov tests, and unpaired Student’s t-test were carried out using IBM SPSS Statistics, while two-way ANOVA with Fisher’s LSD post-hoc tests were performed with STATISTICA. Data were expressed as mean ± standard error of the mean (SEM) Statistical levels of significance are shown in each Fig. as follows: *p < 0.05; **p < 0.01; ***p < 0.001.

## Acknowledgements

We thank the *Caenorhabditis* Genetics Center for *C. elegans* strains, Wormbase for collecting and providing data on *C. elegans*. We are grateful to Somogyvári Milán for technical and methodological help, and other members of the Stress Group for discussions. C.S. is thankful for the Merit Prize of the Semmelweis University.

## Author information

### Authors’ Contributions

CS conceived the study. GH and CS designed the experiments. GH, EG, IT and IM performed the experiments. GH, EG, IT, IM and CS analyzed the data. HG and CS wrote the manuscript with comments from IT. All authors read and approved the manuscript.

## Declarations

### Ethics approval and consent to participate

Not applicable.

### Consent for publication

Not applicable.

### Availability of data and materials

The datasets used and/or analysed during the current study are available from the corresponding author on reasonable request.

## Supplementary Figures

**Fig. S1.**
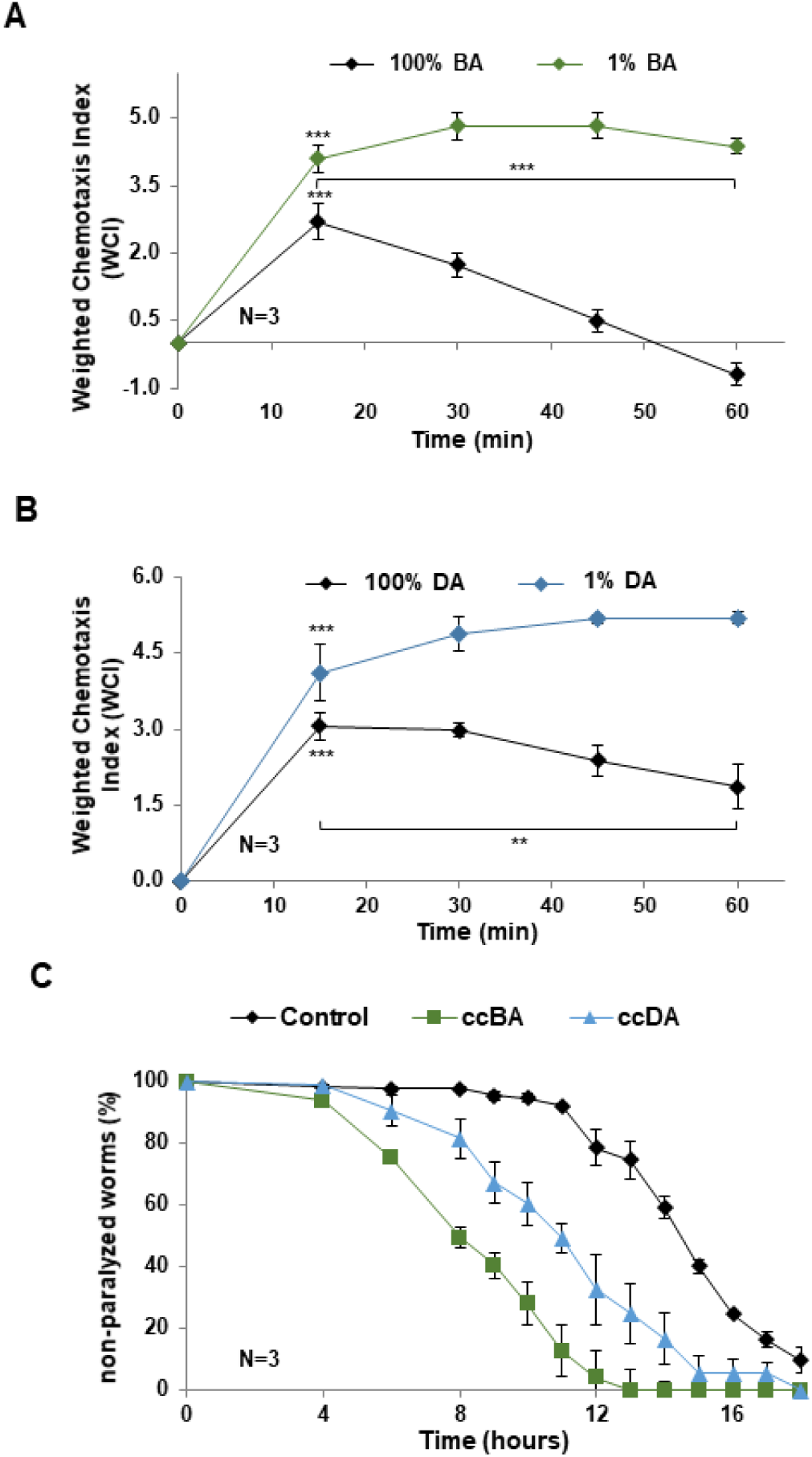
Concentrated benzaldehyde (ccBA) and diacetyl (ccDA) induce behavioral and physiological alterations characteristic to toxicity, Related to Fig. 1. Undiluted, 100% BA and DA trigger initial attraction that diminishes over time as opposed to sustained attraction to diluted 1% odors (A, B). Odor sources were placed in the positive side in three drops (see Methods) opposite to three drops of ethanol vehicle. (C) Continuous exposure to ccBA and ccDA impairs thermotolerance. Error bars represent mean +/− SEM. N = number of independent experiments. *p < 0.05; **p < 0.01; ***p < 0.001. Mean durations of heat shock that induced 50% paralysis by log rank (Mantel-Cox) test were as follows: 14.46 ± 0.23 hours for vehicle treated control, 10.74 ± 0.42 hours for ccBA exposed (p=0.0001 compared to control), 12.45 ± 0.43 hours for ccDA exposed (p=0.011 compared to control).

**Fig. S2.**
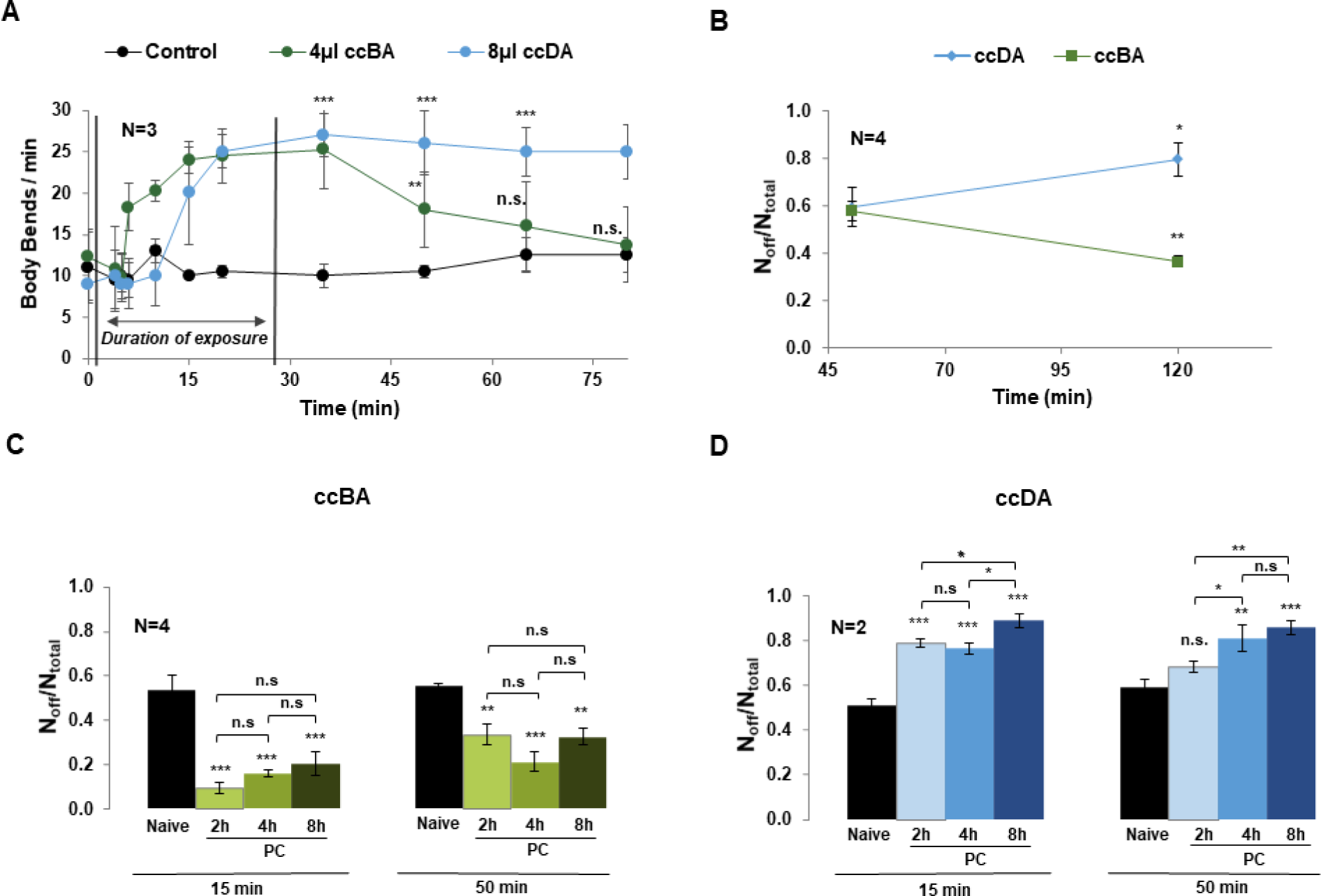
Opposing effects of odor pre-exposure on odor induced behaviors, Related to Fig. 2. (A) Motility assays show a reversible vs. a sustained elevation in locomotion in response to a transient exposure to ccBA *vs.* ccDA. (B) Food aversion data showing that extended odor exposure to ccBA decreases, whereas that to ccDA further increases aversive behavior. (C) ccBA induced food avoidance as a function of duration of preconditioning exposure. (D) ccDA induced food avoidance as a function of duration of preconditioning exposure. Data are expressed as mean ± SEM. N = number of independent experiments. n.s.: not significant, *p < 0.05; **p < 0.01; ***p < 0.001.

**Fig. S3.**
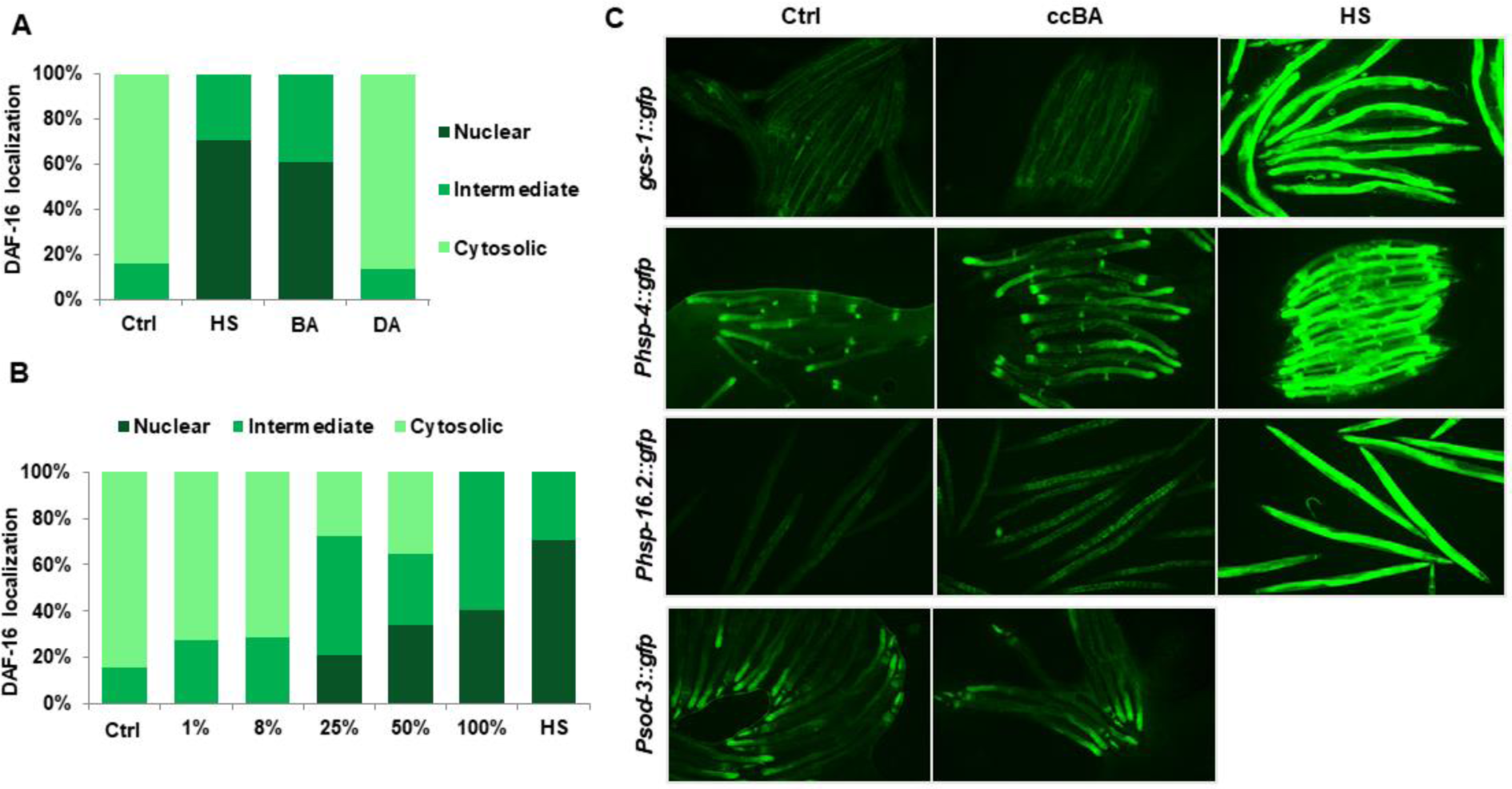
Characteristics of DAF-16 nuclear translocation and specificity of ccBA-induced cytoprotective responses, Related to Fig. 3. (A) ccBA induces nuclear translocation of DAF-16::GFP comparable to that upon heat shock (HS) (quantification of fluorescence intensities of the experiment from Fig. 3A). (B) Concentration dependence of ccBA-induced DAF-16::GFP nuclear translocation. (C) Representative epifluorescent microscopic images of *hsp-16.2::GFP*, *hsp-4::GFP*, *gcs-1::GFP and sod-3::GFP* stress reporters upon ccBA toxicity and heat shock (HS) as a control. Data are expressed as mean of three independent experiments by 40-60 animals evaluated per condition. n.s.: not significant, *p < 0.05; **p < 0.01; ***p < 0.001.

**Fig. S4.**
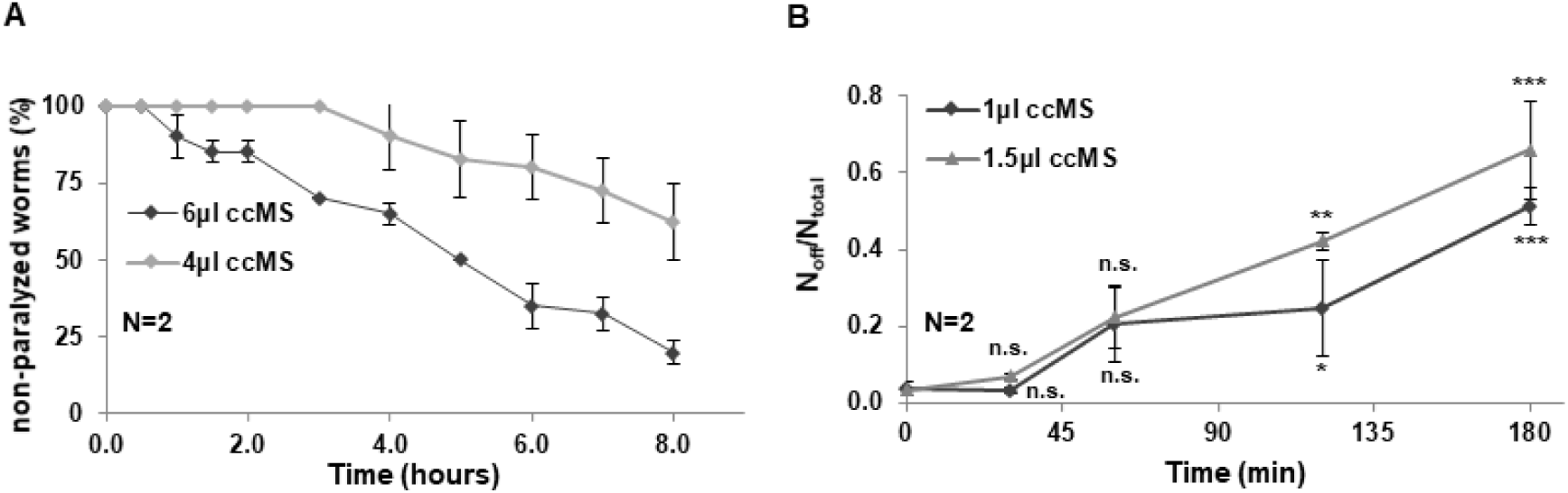
Concentrated methyl-salicylate (ccMS) exposure impairs survival and triggers food avoidance behavior, Related to Fig. 5. (A) Survival curves of nematodes exposed to various doses of ccMS using a hanging drop assay. (B) Time-dependence of ccMS induced food avoidance behavior. Error bars represent mean ± SEM, N = number of independent experiments. *p < 0.05, **p<0.01, ***p<0.001. Mean paralysis values by log rank (Mantel-Cox) test were as follows: 6.65 ± 0.32 hours for 4 μl ccMS, 4.72 ± 0.42 hours for 6 μl ccMS, p=0.001.

## References

1. R. Bijlsma, V. Loeschcke, in Journal of Evolutionary Biology (2005), vol. 18, pp. 744–749.

2. D. Kültz, Molecular and evolutionary basis of the cellular stress response. Annu. Rev. Physiol. 67, 225–57 (2005).

3. H. Selye, The Evolution of the Stress Concept: The originator of the concept traces its development from the discovery in 1936 of the alarm reaction to modern therapeutic applications of syntoxic and catatoxic hormones. Am. Sci. 61, 692–699 (1973).

4. C. J. Kenyon, The genetics of ageing. Nature. 464, 504–512 (2010).

5. A. V Badyaev, Stress-induced variation in evolution: from behavioural plasticity to genetic assimilation. Proc. R. Soc. B Biol. Sci. 272, 877–886 (2005).

6. D. V. Baldwin, Primitive mechanisms of trauma response: An evolutionary perspective on trauma-related disorders. Neurosci. Biobehav. Rev. 37, 1549–1566 (2013).

7. R. McCarty, Learning about stress: neural, endocrine and behavioral adaptations. Stress. 19, 449–475 (2016).

8. E. L. Ardiel, C. H. Rankin, An elegant mind: learning and memory in Caenorhabditis elegans. Learn. Mem. 17, 191–201 (2010).

9. H. C. Barrett, J. Broesch, Prepared social learning about dangerous animals in children. Evol. Hum. Behav. 33, 499–508 (2012).

10. D. Puzzo, J. Fiorito, R. Purgatorio, W. Gulisano, A. Palmeri, O. Arancio, R. Nicholls, Molecular Mechanisms of Learning and Memory. Genes, Environ. Alzheimer’s Dis., 1–27 (2016).

11. R. Garcia, Neurobiology of fear and specific phobias. Learn. Mem. 24, 462–471 (2017).

12. M. Botvinick, T. Braver, Motivation and Cognitive Control: From Behavior to Neural Mechanism. Annu. Rev. Psychol. 66, 83–113 (2015).

13. T. W. Robbins, B. J. Everitt, Neurobehavioural mechanisms of reward and motivation. Curr. Opin. Neurobiol. 6, 228–236 (1996).

14. L. Cosmides, J. Tooby, Evolutionary Psychology: New Perspectives on Cognition and Motivation. Annu. Rev. Psychol. 64, 201–229 (2013).

15. C. I. Bargmann, E. Hartwieg, H. R. Horvitz, Odorant-selective genes and neurons mediate olfaction in C. elegans. Cell. 74, 515–527 (1993).

16. C. Bargmann, Chemosensation in C. elegans. WormBook, 1–29 (2006).

17. O. Hobert, Behavioral plasticity inC. elegans: Paradigms, circuits, genes. J. Neurobiol. 54, 203–223 (2003).

18. T. K. Blackwell, M. J. Steinbaugh, J. M. Hourihan, C. Y. Ewald, M. Isik, SKN-1/Nrf, stress responses, and aging in Caenorhabditis elegans. Free Radic. Biol. Med. 88, 290–301 (2015).

19. S. Ogg, S. Paradis, S. Gottlieb, G. I. Patterson, L. Lee, H. A. Tissenbaum, G. Ruvkun, The Fork head transcription factor DAF-16 transduces insulin-like metabolic and longevity signals in C. elegans. Nature. 389, 994–999 (1997).

20. A. Zečić, B. P. Braeckman, DAF-16/FoxO in Caenorhabditis elegans and Its Role in Metabolic Remodeling. Cells. 9, 109 (2020).

21. A.-L. Hsu, C. T. Murphy, C. Kenyon, Regulation of aging and age-related disease by DAF-16 and heat-shock factor. Science. 300, 1142–5 (2003).

22. S.-K. Park, P. M. Tedesco, T. E. Johnson, Oxidative stress and longevity in Caenorhabditis elegans as mediated by SKN-1. Aging Cell. 8, 258–269 (2009).

23. H. Inoue, N. Hisamoto, J. H. An, R. P. Oliveira, E. Nishida, T. K. Blackwell, K. Matsumoto, The C. elegans p38 MAPK pathway regulates nuclear localization of the transcription factor SKN-1 in oxidative stress response. Genes Dev. 19, 2278–83 (2005).

24. D. Papp, P. Csermely, C. Sőti, A role for SKN-1/Nrf in pathogen resistance and immunosenescence in Caenorhabditis elegans. PLoS Pathog. 8, e1002673 (2012).

25. E. J. Calabrese, K. A. Bachmann, A. J. Bailer, P. M. Bolger, J. Borak, L. Cai, N. Cedergreen, M. G. Cherian, C. C. Chiueh, T. W. Clarkson, R. R. Cook, D. M. Diamond, D. J. Doolittle, M. A. Dorato, S. O. Duke, L. Feinendegen, D. E. Gardner, R. W. Hart, K. L. Hastings, A. W. Hayes, G. R. Hoffmann, J. A. Ives, Z. Jaworowski, T. E. Johnson, W. B. Jonas, N. E. Kaminski, J. G. Keller, J. E. Klaunig, T. B. Knudsen, W. J. Kozumbo, T. Lettieri, S.-Z. Liu, A. Maisseu, K. I. Maynard, E. J. Masoro, R. O. McClellan, H. M. Mehendale, C. Mothersill, D. B. Newlin, H. N. Nigg, F. W. Oehme, R. F. Phalen, M. A. Philbert, S. I. S. Rattan, J. E. Riviere, J. Rodricks, R. M. Sapolsky, B. R. Scott, C. Seymour, D. A. Sinclair, J. Smith-Sonneborn, E. T. Snow, L. Spear, D. E. Stevenson, Y. Thomas, M. Tubiana, G. M. Williams, M. P. Mattson, Biological stress response terminology: Integrating the concepts of adaptive response and preconditioning stress within a hormetic dose–response framework. Toxicol. Appl. Pharmacol. 222, 122–128 (2007).

26. J. A. Melo, G. Ruvkun, Inactivation of Conserved C. elegans Genes Engages Pathogen- and Xenobiotic-Associated Defenses. Cell. 149, 452–466 (2012).

27. J. Singh, A. Aballay, Microbial Colonization Activates an Immune Fight-and-Flight Response via Neuroendocrine Signaling. Dev. Cell. 49, 89–99.e4 (2019).

28. E. Gecse, B. Gilányi, M. Csaba, G. Hajdú, C. Sőti, A cellular defense memory imprinted by early life toxic stress. Sci. Rep. 9, 18935 (2019).

29. W. M. Nuttley, S. Harbinder, D. van der Kooy, Regulation of distinct attractive and aversive mechanisms mediating benzaldehyde chemotaxis in Caenorhabditis elegans. Learn. Mem. 8, 170–81 (2001).

30. K. Yoshida, T. Hirotsu, T. Tagawa, S. Oda, T. Wakabayashi, Y. Iino, T. Ishihara, Odour concentration-dependent olfactory preference change in C. elegans. Nat. Commun. 3, 739 (2012).

31. L. Avery, Y.-J. You, C. elegans feeding. WormBook, 1–23 (2012).

32. T. Tabatabaie, R. A. Floyd, Inactivation of Glutathione Peroxidase by Benzaldehyde. Toxicol. Appl. Pharmacol. 141, 389–393 (1996).

33. A. Andersen, Final report on the safety assessment of benzaldehyde. Int. J. Toxicol. 25 **Suppl 1**, 11–27 (2006).

34. M. Somogyvári, E. Gecse, C. Sőti, DAF-21/Hsp90 is required for C. elegans longevity by ensuring DAF-16/FOXO isoform A function. Sci. Rep. 8, 12048 (2018).

35. K. P. Choe, A. J. Przybysz, K. Strange, The WD40 repeat protein WDR-23 functions with the CUL4/DDB1 ubiquitin ligase to regulate nuclear abundance and activity of SKN-1 in Caenorhabditis elegans. Mol. Cell. Biol. 29, 2704–15 (2009).

36. W. M. Winston, C. Molodowitch, C. P. Hunter, Systemic RNAi in C. elegans requires the putative transmembrane protein SID-1. Science. 295, 2456–9 (2002).

37. A. Galal, L. Walker, I. Khan, Induction of GST and Related Events by Dietary Phytochemicals: Sources, Chemistry, and Possible Contribution to Chemoprevention. Curr. Top. Med. Chem. 14, 2802–2821 (2015).

38. W. P. Jorissen, P. A. A. van der Beek, The oxidation of benzaldehyde. Recl. des Trav. Chim. des Pays-Bas. 49, 138–141 (2010).

39. M. Sankar, E. Nowicka, E. Carter, D. M. Murphy, D. W. Knight, D. Bethell, G. J. Hutchings, The benzaldehyde oxidation paradox explained by the interception of peroxy radical by benzyl alcohol. Nat. Commun. 5, 3332 (2014).

40. S.-W. Park, E. Kaimoyo, D. Kumar, S. Mosher, D. F. Klessig, Methyl salicylate is a critical mobile signal for plant systemic acquired resistance. Science. 318, 113–6 (2007).

41. A. Kauffman, L. Parsons, G. Stein, A. Wills, R. Kaletsky, C. Murphy, C. elegans positive butanone learning, short-term, and long-term associative memory assays. J. Vis. Exp., 1–10 (2011).

42. H. Amano, I. N. Maruyama, Aversive olfactory learning and associative long-term memory in Caenorhabditis elegans. Learn. Mem. 18, 654–65 (2011).

43. E. R. Troemel, J. H. Chou, N. D. Dwyer, H. A. Colbert, C. I. Bargmann, Divergent seven transmembrane receptors are candidate chemosensory receptors in C. elegans. Cell. 83, 207–218 (1995).

44. R. Aoki, T. Yagami, H. Sasakura, K. -i. Ogura, Y. Kajihara, M. Ibi, T. Miyamae, F. Nakamura, T. Asakura, Y. Kanai, Y. Misu, Y. Iino, M. Ezcurra, W. R. Schafer, I. Mori, Y. Goshima, A Seven-Transmembrane Receptor That Mediates Avoidance Response to Dihydrocaffeic Acid, a Water-Soluble Repellent in Caenorhabditis elegans. J. Neurosci. 31, 16603–16610 (2011).

45. W. M. Kluwe, C. A. Montgomery, H. D. Giles, J. D. Prejean, Encephalopathy in rats and nephropathy in rats and mice after subchronic oral exposure to benzaldehyde. Food Chem. Toxicol. 21, 245–250 (1983).

46. J. M. McKim, P. K. Schmieder, G. J. Niemi, R. W. Carlson, T. R. Henry, Use of respiratory-cardiovascular responses of rainbow trout (*Salmo gairdneri*) in identifying acute toxicity syndromes in fish: Part 2. malathion, carbaryl, acrolein and benzaldehyde. Environ. Toxicol. Chem. 6, 313–328 (1987).

47. D. M. Brass, S. M. Palmer, Models of toxicity of diacetyl and alternative diones. Toxicology. 388, 15–20 (2017).

48. S. Clark, C. K. Winter, Diacetyl in Foods: A Review of Safety and Sensory Characteristics. Compr. Rev. Food Sci. Food Saf. 14, 634–643 (2015).

49. T. Shibamoto, Diacetyl: Occurrence, Analysis, and Toxicity. J. Agric. Food Chem. 62, 4048–4053 (2014).

50. W. M. Nuttley, K. P. Atkinson-Leadbeater, D. Van Der Kooy, Serotonin mediates food-odor associative learning in the nematode Caenorhabditiselegans. Proc. Natl. Acad. Sci. U. S. A. 99, 12449–54 (2002).

51. H. Sasakura, I. Mori, Behavioral plasticity, learning, and memory in C. elegans. Curr. Opin. Neurobiol. 23, 92–99 (2013).

52. V. Singh, A. Aballay, Heat-shock transcription factor (HSF)-1 pathway required for Caenorhabditis elegans immunity. Proc. Natl. Acad. Sci. U. S. A. 103, 13092–7 (2006).

53. J. M. A. Tullet, M. Hertweck, J. H. An, J. Baker, J. Y. Hwang, S. Liu, R. P. Oliveira, R. Baumeister, T. K. Blackwell, Direct Inhibition of the Longevity-Promoting Factor SKN-1 by Insulin-like Signaling in C. elegans. Cell. 132, 1025–1038 (2008).

54. T. Greene, S. Rogers, A. Franzen, R. Gentry, A critical review of the literature to conduct a toxicity assessment for oral exposure to methyl salicylate. Crit. Rev. Toxicol. 47, 98–120 (2017).

55. J. M. Kyriakis, J. Avruch, Mammalian MAPK Signal Transduction Pathways Activated by Stress and Inflammation: A 10-Year Update. Physiol. Rev. 92, 689–737 (2012).

56. J. A. Melo, G. Ruvkun, Inactivation of conserved C. elegans genes engages pathogen- and xenobiotic-associated defenses. Cell. 149, 452–66 (2012).

57. D. H. Kim, R. Feinbaum, G. Alloing, F. E. Emerson, D. A. Garsin, H. Inoue, M. Tanaka-Hino, N. Hisamoto, K. Matsumoto, M.-W. Tan, F. M. Ausubel, A conserved p38 MAP kinase pathway in Caenorhabditis elegans innate immunity. Science. 297, 623–6 (2002).

58. K. Lee, E. Mylonakis, An Intestine-Derived Neuropeptide Controls Avoidance Behavior in Caenorhabditis elegans. Cell Rep. 20, 2501–2512 (2017).

59. H. Xiong, C. Pears, A. Woollard, An enhanced C. elegans based platform for toxicity assessment. Sci. Rep. 7, 9839 (2017).

60. A. M. Horspool, H. C. Chang, Neuron-specific regulation of superoxide dismutase amid pathogen-induced gut dysbiosis. Redox Biol. 17, 377–385 (2018).

61. A. Livnat, Interaction-based evolution: how natural selection and nonrandom mutation work together. Biol. Direct. 8, 24 (2013).

62. J. Luo, Z. Xu, Z. Tan, Z. Zhang, L. Ma, Neuropeptide receptors NPR-1 and NPR-2 regulate Caenorhabditis elegans avoidance response to the plant stress hormone methyl salicylate. Genetics. 199, 523–31 (2015).

63. Y. Zhang, H. Lu, C. I. Bargmann, Pathogenic bacteria induce aversive olfactory learning in Caenorhabditis elegans. Nature. 438, 179–184 (2005).

64. R. Nakad, L. B. Snoek, W. Yang, S. Ellendt, F. Schneider, T. G. Mohr, L. Rösingh, A. C. Masche, P. C. Rosenstiel, K. Dierking, J. E. Kammenga, H. Schulenburg, Contrasting invertebrate immune defense behaviors caused by a single gene, the Caenorhabditis elegans neuropeptide receptor gene npr-1. BMC Genomics. 17, 280 (2016).

65. G. Harris, T. Wu, G. Linfield, M.-K. Choi, H. Liu, Y. Zhang, Molecular and cellular modulators for multisensory integration in C. elegans. PLOS Genet. 15, e1007706 (2019).

66. J. Singh, A. Aballay, Intestinal infection regulates behavior and learning via neuroendocrine signaling. Elife. 8 (2019), doi:10.7554/eLife.50033.

67. G. M. Stein, C. T. Murphy, C. elegans positive olfactory associative memory is a molecularly conserved behavioral paradigm. Neurobiol. Learn. Mem. 115, 86–94 (2014).

68. W. B. Cannon, The wisdom of the body, 2nd ed. (1939).

69. A. Pittig, M. Treanor, R. T. LeBeau, M. G. Craske, The role of associative fear and avoidance learning in anxiety disorders: Gaps and directions for future research. Neurosci. Biobehav. Rev. 88, 117–140 (2018).

70. S. Brenner, The genetics of Caenorhabditis elegans. Genetics. 77, 71–94 (1974).

71. N. A. Croll, Components and patterns in the behaviour of the nematode Caenorhabditis elegans. J. Zool. 176, 159–176 (2009).

72. M. D. Gyurkó, P. Csermely, C. Sőti, A. Steták, Distinct roles of the RasGAP family proteins in C. elegans associative learning and memory. Sci. Rep. 5, 15084 (2015).

73. K. Milward, K. E. Busch, R. J. Murphy, M. de Bono, B. Olofsson, Neuronal and molecular substrates for optimal foraging in Caenorhabditis elegans. Proc. Natl. Acad. Sci. U. S. A. 108, 20672–7 (2011).

74. E. R. Sawin, R. Ranganathan, H. R. Horvitz, C. elegans Locomotory Rate Is Modulated by the Environment through a Dopaminergic Pathway and by Experience through a Serotonergic Pathway. Neuron. 26, 619–631 (2000).

75. L. Timmons, in C. elegans (Humana Press, New Jersey, 2006; http://www.ncbi.nlm.nih.gov/pubmed/16988430), vol. 351, pp. 119–126.

